# Elastic properties and shape of the Piezo dome underlying its mechanosensory function

**DOI:** 10.1101/2022.07.01.498002

**Authors:** Christoph A. Haselwandter, Yusong R. Guo, Ziao Fu, Roderick MacKinnon

**Author notes:** Christoph A. Haselwandter and Roderick MacKinnon, **Email:** (CAH); (RM). These authors contributed equally to this work (Christoph A. Haselwandter, Roderick MacKinnon). **Author Contributions:** RM and YRG and ZF designed the experiments. YRG and ZF expressed and purified Piezo1 protein, reconstituted Piezo vesicles, collected tomograms, processed images and digitized the oriented Piezo vesicle images. CAH and RM conceived the project, developed the theory of Piezo vesicle mechanics, analyzed the data beginning with the oriented Piezo vesicle images and wrote the paper. **Competing Interest Statement:** The authors declare no competing interests.

## Abstract

In a companion paper we show that the free membrane shape of lipid bilayer vesicles containing the mechanosensitive ion channel Piezo can be predicted, with no free parameters, from membrane elasticity theory together with measurements of the protein geometry and vesicle size (<u>accompanying paper</u>). Here we use these results to determine the force that Piezo exerts on the free membrane and, hence, that the free membrane exerts on Piezo, for a range of vesicle sizes. From vesicle shape measurements alone, we thus obtain a force-distortion relationship for Piezo, from which we deduce Piezo’s intrinsic radius of curvature, 42 ± 12 nm, and bending stiffness, 18 ± 2.1 *k*_*B*_ *T*, in free-standing lipid bilayer membranes mimicking cell membranes. Applying these estimates to a spherical cap model of Piezo embedded in a lipid bilayer, we suggest that Piezo’s intrinsic curvature, surrounding membrane footprint, small stiffness, and large area are the key properties of Piezo that give rise to low-threshold, high-sensitivity mechanical gating.

## Introduction

Piezo 1 and 2 are called mechanosensitive ion channels because they conduct ions across the cell membrane when a mechanical force is applied to the cell(1). Thus, Piezo channels are somehow rigged to enable a mechanical force to open their pore. The ensuing ion conduction triggers subsequent processes inside the cell, culminating in a cell’s response to the mechanical force. This sequence of events, force on the membrane → ion conduction → cell response, is central to numerous biological processes(2, 3). Mediated by Piezo channels, these include, but are not limited to, volume regulation in red blood cells, the control of vascular blood pressure, and the sensation of touch(4, 5).

What physical properties endow Piezo channels with responsiveness to mechanical force? Their unique shape among ion channels has inspired one proposal, known as the ‘membrane dome model’(6). Piezo channels in their closed conformation are curved, in contrast to most other ion channels and membrane proteins, which exhibit an approximately planar arrangement of transmembrane helices(7). Consequently, Piezo channels locally curve the membrane into a ‘Piezo dome’ and surrounding ‘membrane footprint’(8). The membrane dome model posits that an open Piezo channel will be less curved, more like other membrane proteins(6, 8, 9), and it is known that Piezo can change its shape. Cryo-electron microscopy (cryo-EM) studies have shown that Piezo channels in lipid bilayer vesicles change their curvature depending on vesicle size(<u>accompanying paper</u>(9)). High-speed atomic force microscopy (HS-AFM) has been used to flatten Piezo(9) and, by cryo-EM, nearly flat Piezo channels with a pore that appears to be somewhat widened compared to curved Piezo channels have been observed(10). If it is true that Piezo must reduce its curvature to open its pore, then increased lateral membrane tension should favor the open conformation by a work energy term *γ* Δ*A*_proj_, where *γ* is the lateral membrane tension and Δ*A*_proj_ is the expansion of Piezo’s projected, in-plane area due to its reduced curvature, which includes contributions arising from Piezo’s membrane footprint(6, 8). The membrane dome model is still unproven, but it rationalizes Piezo’s highly unusual, curved shape.

Beyond shape alone, to understand Piezo’s responsiveness to mechanical force, we need to know how its shape changes when force is applied. To use an analogy: to describe a spring’s mechanical properties we need to know how its shape, that is its length, depends on force; for a linear spring, for instance, the spring’s length is proportional to load, and this dependence is captured in the Hooke constant (spring stiffness). Structural biology lets us determine a molecule’s shape, but how can we do this while applying a force to a molecule? Furthermore, how do we apply a force while the molecule, in this case a mechanosensitive ion channel, resides in an unsupported, free membrane environment?

When a Piezo ion channel resides in a lipid bilayer vesicle, the curved lipid membrane surrounding the channel exerts a force on it. Different sized vesicles exert different values of force, causing the channel to change its shape. In a companion paper, we developed a continuum elasticity theory of Piezo vesicle shape and showed that this theory predicts Piezo vesicle shapes that are in quantitative agreement with those observed experimentally (<u>accompanying paper</u>). In this study, we build on this work to deduce Piezo’s force-shape relationship, which we then relate to Piezo’s mechanosensitive gating. In particular, we extract the force curve associated with the Piezo vesicles described in our companion paper, and apply it to understand how Piezo’s curvature, stiffness, and area give rise to its mechanosensory properties.

A point regarding notation: As in our companion paper, we refer with the term ‘Piezo channel’ to the ion channel protein. ‘Piezo dome’ refers to the functional unit consisting of the ion channel protein plus the lipid bilayer contained within the channel’s approximate perimeter, with the lipid membrane connecting smoothly across the Piezo dome boundary. We refer to the region of free membrane, or free lipid bilayer, outside the Piezo dome that is deformed by Piezo as Piezo’s ‘membrane footprint.’ ‘Piezo vesicle’ refers to a lipid bilayer vesicle containing Piezo.

## Results

### Shape of Piezo vesicles

Before deriving the force curve of the Piezo dome, we review some of the results of our companion paper most pertinent to the present study. In our companion paper we show how membrane elasticity theory can be used to predict the shape of the free membrane, outside the Piezo dome, in Piezo vesicles (<u>accompanying paper</u>). Our starting point is thereby the Helfrich energy equation for the free membrane,

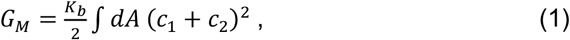

where the constant *K*_*b*_ is the lipid bilayer bending modulus, *c*_1_ and *c*_2_ are the principal curvatures of the mid-membrane surface, and the integral is carried out over the entire free membrane surface (11). We use *K*_*b*_ = 20 *k*_*B*_*T*, which approximates the bending modulus of the lipid bilayers in our Piezo vesicles(12). Through Eq. 1, each possible free membrane shape gives an associated value of *G*_*M*_. We determine the free membrane shape minimizing *G*_*M*_ by solving the Hamilton equations associated with Eq. 1, subject to suitable constraints. One key constraint thereby specifies the size of Piezo vesicles. We find it convenient to define the vesicle size through the radius of a hypothetical sphere comprising the Piezo dome plus the free membrane, *R*_*ν*_ (<u>accompanying paper</u>). Furthermore, Eq. 1 suggests that two key properties of the Piezo dome—the Piezo dome-free membrane contact angle, *α*, and the projected, in-plane radius of the Piezo dome, *r*_*b*_—govern how Piezo affects the free membrane shape (see Fig. 1). Comparing the free membrane shapes calculated from Eq. 1 for seven Piezo vesicles, ranging in size from *R*_*ν*_ ≈ 12.1 nm to *R*_*ν*_ ≈ 36.2 nm, to the corresponding shapes obtained by tomographic reconstruction of cryo-EM images, we find that minimization of Eq. 1 successfully predicts the free membrane shape of Piezo vesicles, without any free parameters (Fig. 1).

**Fig. 1:**
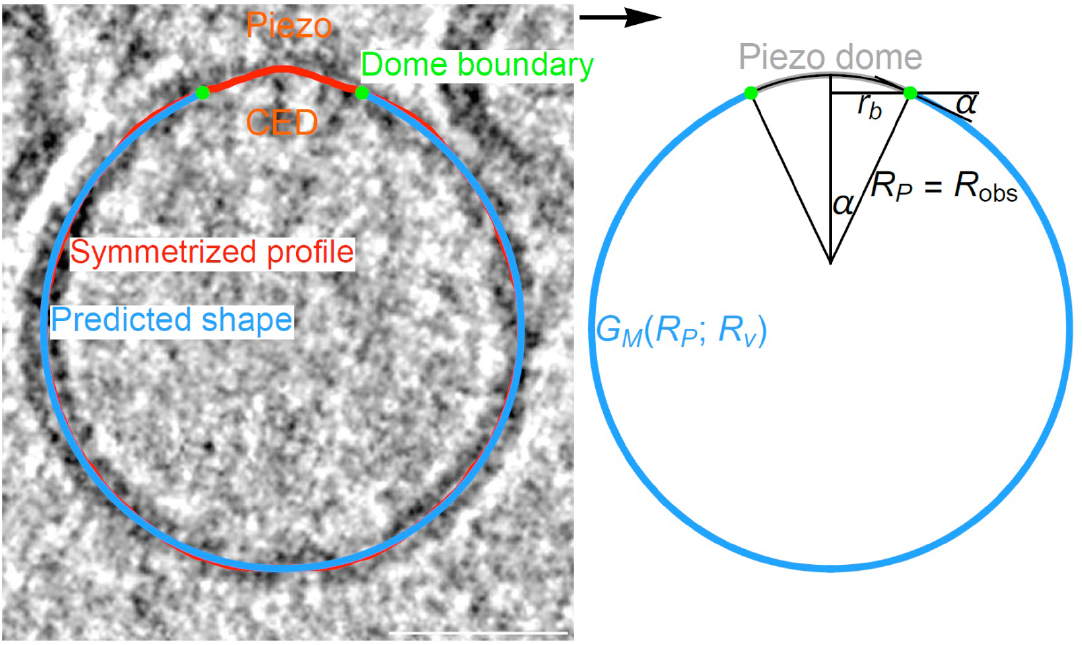
Predicting the shape of Piezo vesicles. Oriented Piezo vesicle image obtained by cryo-EM tomography (left panel) and associated symmetrized (measured) Piezo vesicle profile (red curve), Piezo dome boundary obtained by integrating out a vesicle surface area equal to *A*_*P*_ = 450 nm^2^ starting at the vesicle north pole (green dots), and corresponding predicted free membrane profile (blue curves). The predicted Piezo vesicle profile is obtained, with no free parameters, from the membrane elasticity theory of Piezo vesicle shape described in our companion paper (<u>accompanying paper</u>), and minimizes the Helfrich energy equation, Eq. 1. The value of the minimized free membrane bending energy, *G*_*M*_(*R*_*P*_; *R*_*ν*_) in Eq. 1, depends on the vesicle size *R*_*ν*_ and on the radius of curvature at the Piezo dome boundary, *R*_*P*_, with *R*_obs_ denoting the value of *R*_*P*_ associated with the predicted free membrane shapes in Fig. 4 of our companion paper. We obtain *R*_obs_ from 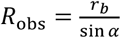, where *r*_*b*_ and *α* denote the in-plane Piezo dome radius and the Piezo dome contact angle associated with the predicted free membrane shapes in Fig. 4 of our companion paper, respectively. The vesicle shown here corresponds to vesicle 6 with *R*_*ν*_ ≈ 35.0 nm in Fig. 4 of our companion paper. Scale bar, 26 nm.

The shape of the Piezo dome depends on the Piezo vesicle radius *R*_*ν*_, with the Piezo dome becoming less curved as *R*_*ν*_ is increased. In our companion paper we show that, if one models the Piezo dome as a spherical cap with fixed area *A*_cap_, the measured changes in the shape of the Piezo dome can, approximately, be captured by a single parameter, the Piezo dome radius of curvature 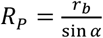 (Fig. 1) (<u>accompanying paper</u>). For a given value of *A*_cap_, this model of the Piezo dome geometrically defines the Piezo dome properties affecting membrane shape deformations in Eq. 1, *α* and *r*_*b*_, as a function of *R*_*P*_(13). For each measured Piezo vesicle, we choose here *A*_cap_ so that we obtain the values of *α* and *r*_*b*_ associated with the predicted free membrane shapes in Fig. 4 of our companion paper, and denote the corresponding value of *R*_*P*_ by *R*_obs_. To be clear, our approach for calculating the forces in Piezo vesicles does not rely on the spherical cap model of the Piezo dome. We employ here this model because it is simple and seems to capture, for the Piezo dome, the approximate relation between *α* and *r*_*b*_.

### The force curve

As reviewed above, our companion paper demonstrates that minimization of the Helfrich energy equation, Eq. 1, successfully predicts the observed shapes of the free membrane in Piezo vesicles, over a range of vesicle radii *R*_*ν*_ (<u>accompanying paper</u>). We denote this minimum energy by *G*_*M*_(*R*_*P*_; *R*_*ν*_), to emphasize that distinct vesicle sizes *R*_*ν*_ yield, in general, a distinct dependence of *G*_*M*_ on the Piezo dome radius of curvature *R*_*P*_. We now analyze the balance of forces between the free membrane and the Piezo dome in Piezo vesicles. To demonstrate how this will work, we consider the following thought experiment.

A hypothetical Piezo vesicle, with *R*_*ν*_ ≈ 24.7 nm, contains a Piezo dome that we model, as described above, as a spherical cap with fixed area *A*_cap_ and variable radius of curvature *R*_*P*_ (see Fig. 2*A*). Now imagine that *R*_*P*_ can be adjusted to any desired value; because *A*_cap_ is fixed^1^, *R*_*P*_ will specify *α* and *r*_*b*_. For a given *R*_*P*_, we calculate the free membrane shape by minimizing the Helfrich energy equation, Eq. 1, employing, as input parameter values, the values of *α* and *r*_*b*_ obtained from *R*_*P*_ along with the measured free membrane area. We then enter this shape into Eq. 1 to calculate the corresponding value of the free membrane energy, *G*_*M*_(*R*_*P*_; *R*_*ν*_). Repeating this calculation for a range of *R*_*P*_ values, we graph *G*_*M*_ as a function of *R*_*P*_, keeping *R*_*ν*_ fixed: this graph shows how the free membrane energy varies with *R*_*P*_ for a given Piezo vesicle (Fig. 2*A*). We find that, when Piezo has a radius of curvature 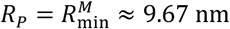, the free membrane energy is least for the Piezo vesicle considered here. Some examples of the predicted free membrane shapes for specific values of *R*_*P*_ are shown (Fig. 2*A*).

**Fig. 2:**
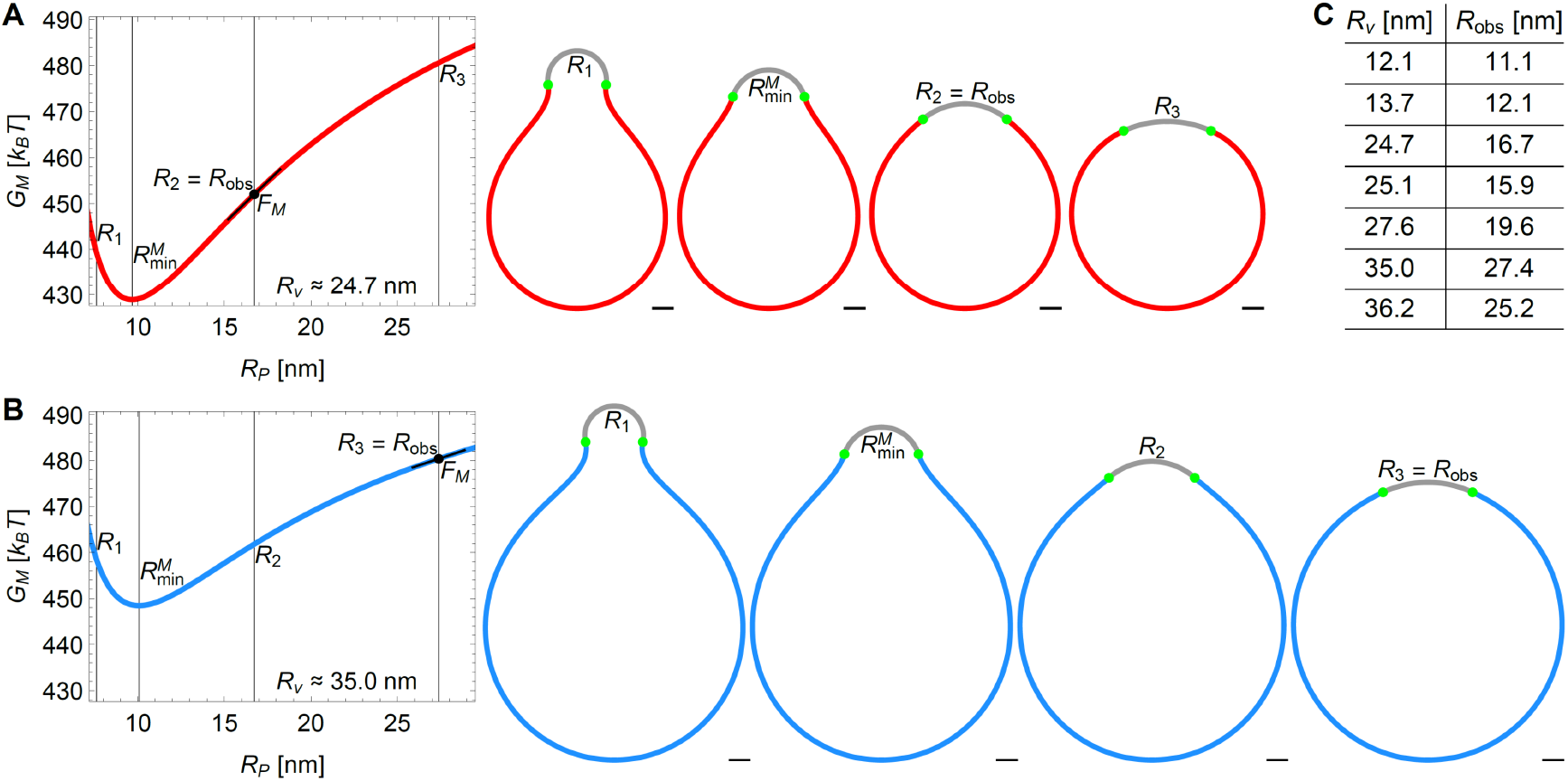
Energy landscape of Piezo vesicle shape. Stationary lipid membrane bending energy *G*_*M*_(*R*_*P*_; *R*_*ν*_) as a function of Piezo dome radius of curvature *R*_*P*_ for (*A*) the Piezo vesicle radius *R*_*ν*_ ≈ 24.7 nm (vesicle 3 in Fig. 4 of our companion paper (<u>accompanying paper</u>)) and (*B*) the Piezo vesicle radius *R*_*ν*_ ≈ 35.0 nm (vesicle 6 in Fig. 4 of our companion paper) together with selected vesicle cross sections at *R*_*P*_ = *R*_1_ = 7.6 nm, the values of *R*_*P*_ minimizing 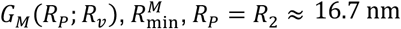, and *R*_*P*_ = *R*_3_ ≈ 27.4 nm (red and blue curves). We have 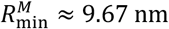 in panel *A* and 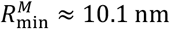 in panel *B*. In panel *A, R*_2_ is equal to the observed value of *R*_*P*_, *R*_2_ = *R*_obs_, while *R*_3_ = *R*_obs_ in panel *B*. For each Piezo vesicle, we describe the Piezo dome geometry as a spherical cap with fixed cap area *A*_cap_. The in-plane Piezo dome radius *r*_*b*_ and the Piezo dome contact angle *α* associated with the predicted free membrane shapes in Fig. 4 of our companion paper determine, for each Piezo vesicle, *A*_cap_ via 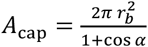. Scale bars, 5 nm. (*C*) Table showing the Piezo dome radius of curvature at the Piezo dome boundary, *R*_*P*_ = *R*_obs_, for the predicted free membrane shapes in Fig. 4 of our companion paper versus *R*_*ν*_. The corresponding spherical cap areas are given by *A*_cap_ ≈ 471 nm^2^, 414 nm^2^, 421 nm^2^ (panel *A*), 471 nm^2^, 428 nm^2^, 444 nm^2^ (panel *B*), and 442 nm^2^ (top to bottom).

The vesicle radius *R*_*ν*_ ≈ 24.7 nm in the above thought experiment corresponds to the radius of measured Piezo vesicle 3 in Fig. 4 of our companion paper. Notably, Piezo vesicle 3, with an observed Piezo dome radius of curvature *R*_*P*_ = *R*_obs_ ≈ 16.7 nm, does not look like the vesicle with least free membrane energy in the thought experiment, which would be associated with a Piezo dome radius of curvature 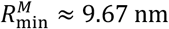 (Fig. 2*A*). The discrepancy between the thought experiment and the measured Piezo vesicle is explicable: the thought experiment calculates only the free membrane contribution to the shape energy of a Piezo vesicle. In other words, if the Piezo dome did not contribute to the total energy of the Piezo vesicle, then it would adopt *R*_*P*_ ≈ 9.67 nm in a Piezo vesicle with *R*_*ν*_ ≈ 24.7 nm. But the Piezo dome clearly does contribute to the total energy of a Piezo vesicle, expressed as *G*_tot_ = *G*_*M*_ + *G*_*P*_, where *G*_*P*_(*R*_*P*_) is the Piezo dome contribution. The minimum energy shape, i.e., the shape corresponding to *R*_*P*_ = *R*_obs_, then must occur when 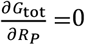. Here, the partial derivative notation reminds us that *G*_tot_ is also a function of the Piezo vesicle radius, *R*_*ν*_, which is held constant in the derivative with respect to *R*_*P*_. From *G*_tot_ = *G*_*M*_ + *G*_*P*_, we have 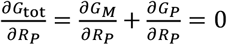 at *R*_*P*_ = *R*_obs_, and therefore 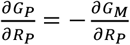 at *R*_*P*_ = *R*_obs_. This last equation, rewritten as

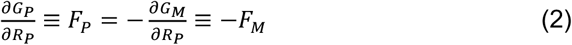

at *R*_*P*_ = *R*_obs_, expresses the equilibrium balance of forces between the Piezo dome and the free membrane in Piezo vesicles. In words, the slope of *G*_*M*_(*R*_*P*_; *R*_*ν*_) with respect to *R*_*P*_, evaluated at *R*_*P*_ = *R*_obs_, equals the force exerted on the free membrane by the Piezo dome, *F*_*M*_, and minus this slope is the force exerted on the Piezo dome by the free membrane, *F*_*P*_. An intuitive understanding of this balance of forces can be grasped through inspection of the hypothetical vesicle shapes for *R*_*ν*_ ≈ 24.7 nm (Fig. 2*A*). The free membrane tends towards its minimum energy shape associated with 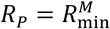, while the Piezo dome tends towards a larger *R*_*P*_, i.e., a flatter shape. A balance of forces is reached at an intermediate radius of curvature, when *R*_*P*_ = *R*_obs_. Thus, the Piezo vesicle’s curved free membrane ‘squeezes’ the Piezo dome, like compressing a spring, to be more curved than it would be in a planar membrane. The force that the free membrane exerts on the Piezo dome in the measured Piezo vesicle 3, i.e., minus the slope of *G*_*M*_(*R*_*P*_; *R*_*ν*_) with respect to *R*_*P*_ at *R*_*P*_ = *R*_obs_, is given by 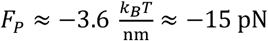. The minus sign here indicates that *F*_*P*_ tends to decrease the value of *R*_*P*_. From this analysis of the measured Piezo vesicle 3, we conclude that when the Piezo dome has a compressive force *F*_*P*_ ≈ −15 pN applied to it by the free vesicle membrane, it has a radius of curvature *R*_*P*_ ≈ 16.7 nm.

In Fig. 2*B* we analyze in the same manner a larger vesicle with radius *R*_*ν*_ ≈ 35.0 nm, which corresponds to the measured Piezo vesicle 6 in Fig. 4 of our companion paper (see also Fig. 1) (<u>accompanying paper</u>). For this vesicle, the experimentally determined *R*_obs_ ≈ 27.4 nm. From Eq. 2, the associated slope of *G*_*M*_(*R*_*P*_; *R*_*ν*_) with respect to *R*_*P*_ yields a force on the Piezo dome *F*_*P*_ ≈ −5.1 pN. Again, the minus sign indicates a compressive force on the Piezo dome by the free vesicle membrane, but in this case the force has a smaller magnitude. Application of Eq. 2 to all seven measured Piezo vesicles (see Fig. 2*C*) yields a set of seven force-displacement values, (*R*_*P*_, *F*_*P*_), graphed in Fig. 3. The graph shows that highly curved vesicles, i.e., vesicles with small *R*_*ν*_, apply a large compressive force to curve the Piezo dome, and that larger vesicles exert a smaller force. Across the range of Piezo vesicle sizes considered here, the force on the Piezo dome ranges from approximately −71 pN to −5.1 pN. In the limit of an infinitely large vesicle, i.e., in a planar membrane, without lateral tension, the surrounding membrane would exert no net elastic force on the Piezo dome. Thus, in a planar, tensionless membrane, the Piezo dome would adopt a shape that conforms to its intrinsic curvature, which we describe next.

**Fig. 3:**
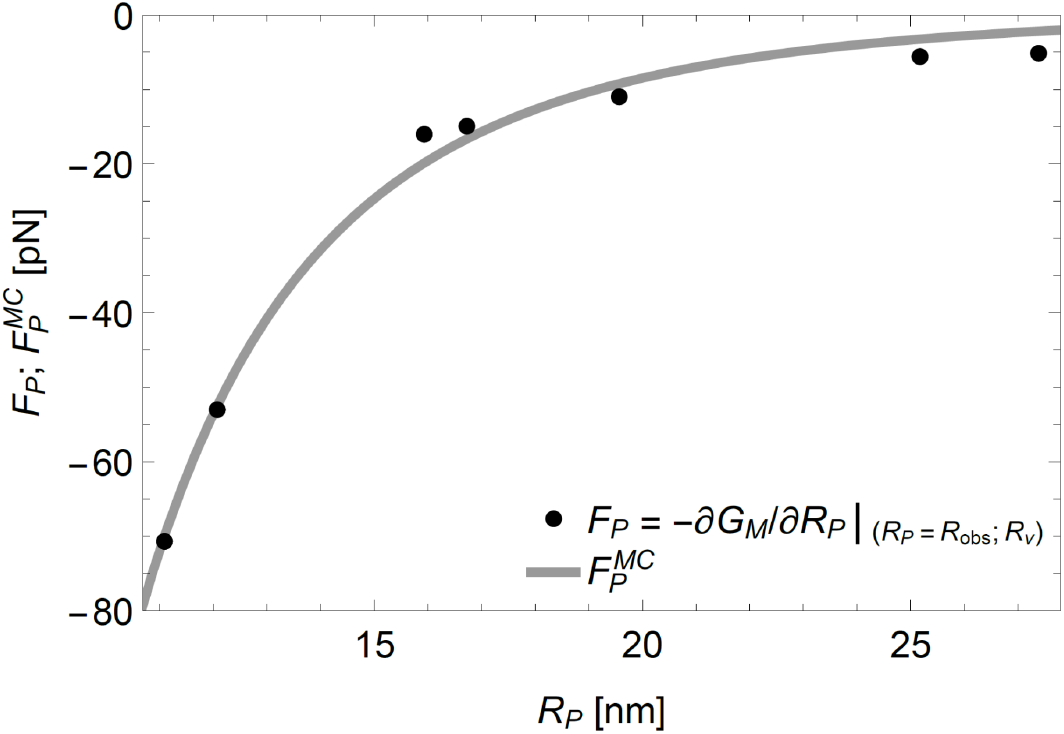
Mechanics of the Piezo dome. Force exerted on the Piezo dome, *F*_*P*_, for the observed Piezo dome radii of curvature, *R*_*P*_ = *R*_obs_, obtained from (minus) the derivative of the stationary free membrane bending energy with respect to 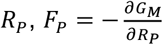 in Eq. 2, at *R*_*P*_ = *R*_obs_ for the measured Piezo vesicle radii *R*_*ν*_ (see also Fig. 2*C*). In mechanical equilibrium of the Piezo-membrane system, the restoring force generated internally by the Piezo dome is given by −*F*_*P*_. The grey curve shows the force on the Piezo dome obtained by fitting the mean curvature force in Eq. 4, 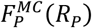, to the force-displacement values of the Piezo dome, (*R*_*P*_, *F*_*P*_). We have the fits *K*_*P*_ = 18 ± 2.1 *k*_*B*_*T* and *R*_0_ = 42 ± 12 nm in Eqs. 3 and 4.

### Relating mechanics to Piezo structure

The force-displacement curve of a spring can be used to deduce key mechanical properties of the spring, such as its stiffness. Similarly, the force-displacement values of the Piezo dome, (*R*_*P*_, *F*_*P*_) in Fig. 3, can be used to deduce mechanical properties of the Piezo dome. To this end, it is instructive to consider, inspired by the Piezo dome shapes found in our companion paper, a highly simplified model of the energetics of the Piezo dome (<u>accompanying paper</u>). In this model, we describe the Piezo dome as a spherical cap of fixed area *A*_*P*_ = 450 nm^2^. Furthermore, we adapt Eq. 1 to the Piezo dome itself, replacing the lipid bilayer bending modulus *K*_*b*_ by the Piezo dome bending modulus *K*_*P*_ and allowing for a preferred, intrinsic radius of curvature of the Piezo dome, *R*_0_. We thus have the following mean curvature energy of the Piezo dome,

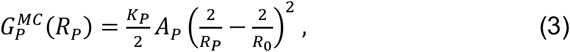

where we have noted that the principal curvatures 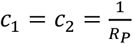 for a spherical cap with radius of curvature *R*_*P*_. According to Eq. 3, the Piezo dome will adopt a radius of curvature *R*_*P*_ = *R*_0_ if it is not perturbed by external forces. In other words, Eq. 3 implies that the Piezo dome’s minimum energy state occurs when *R*_*P*_ = *R*_0_, in analogy to the natural length of a spring that can be compressed or stretched. Differentiating Eq. 3 with respect to *R*_*P*_ gives the mean curvature force,

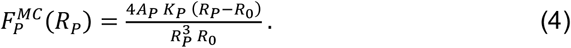

The solid curve in Fig. 3 is a fit of 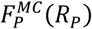 to the force-displacement values of the Piezo dome, (*R*_*P*_, *F*_*P*_). There are two fitting parameters in Eq. 4, *K*_*P*_ (= 18 ± 2.1 *k*_*B*_*T*) and *R*_0_ (= 42 ± 12 nm). The Piezo force curve data in Fig. 3 and, hence, the values of *K*_*P*_ and *R*_0_ depend on the lipid bilayer composition through, for instance, the bilayer bending modulus *K*_*b*_ in Eq. 1 (12).To examine the dependence of *K*_*P*_ and *R*_0_ on the cap area *A*_*P*_, we obtain *K*_*P*_ = 17 ± 2.1 *k*_*B*_*T* and *R*_0_ = 40 ± 11 nm for *A*_*P*_ = 410 nm^2^ and *K*_*P*_ = 18 ± 2.0 *k*_*B*_*T* and *R*_0_ = 45 ± 13 nm for *A*_*P*_ = 490 nm^2^ (see Fig. S1 in *Supplementary Information* section S1). Clearly, the values of *K*_*P*_ and *R*_0_ depend only weakly on the value of *A*_*P*_. This means that uncertainty in the location of the Piezo dome boundary will have little impact on the estimates of *K*_*P*_ and *R*_0_ for the Piezo dome.

While Eqs. 3 and 4 are seen to capture the basic trends in the Piezo force curve data in Fig. 3, we note several potential shortcomings of this model. First, while we provide evidence in a companion paper that the shape of the Piezo dome approximately conforms to a spherical cap geometry, the observed Piezo dome shapes do not, strictly speaking, show constant curvatures, as assumed in Eqs. 3 and 4 (<u>accompanying paper</u>). Second, Eqs. 3 and 4 describe the Piezo dome as a homogeneous material, with constant *K*_*P*_. Since the Piezo dome is composed of both a lipid bilayer and the Piezo protein, this assumption can only be correct in an approximate sense. Third, we focused in Eqs. 3 and 4 on contributions to the Piezo dome mechanics due to the mean curvature of the Piezo dome at the mid-membrane surface and neglected any contributions due to the Gaussian curvature of the Piezo dome. The Gaussian curvature is expected to yield a contribution to the Piezo dome energy that depends on the detailed shape of the bilayer-Piezo boundary, and is unlikely to show the simple dependence on *R*_*P*_ assumed in Eq. 3 (14). Given the limited Piezo force curve data currently available, we focus here on the highly simplified model in Eqs. 3 and 4, which already provides a reasonably good fit with only two fitting parameters.

The force curve in Fig. 3 suggests that the Piezo dome has a radius of curvature *R*_*P*_∼40 nm in a planar membrane without lateral tension, such as one might find on the surface of a mechanically unperturbed cell. The highly curved structures (*R*_*P*_∼10 nm) of Piezo 1 and 2, determined in detergent micelles, indicate that Piezo proteins that are removed from the membrane are most stable in a conformation with high curvature. But, considering the space between Piezo’s extended protein arms, the area of a Piezo dome in a membrane comprises approximately 75% lipid bilayer and 25% protein. As per Eq. 1, the energy cost to curve this bilayer away from a plane is considerable. It seems likely that the increased radius of curvature, from *R*_*P*_∼10 nm observed in isolated Piezo proteins to *R*_*P*_∼40 nm in a planar membrane, reflects an equilibrium balance of forces between the Piezo protein and the bilayer within the Piezo dome (see Fig. S2 in *Supplementary Information* section S1). Whatever the detailed molecular mechanisms involved, the force curve informs us that Piezo in planar cell membranes should be flatter in shape than the highly curved Piezo proteins in detergent micelles.

The bending modulus of the Piezo dome, *K*_*P*_ = 18 ± 2.1 *k*_*B*_*T*, is indistinguishable from the bending modulus of the pure lipid bilayer membrane used here, *K*_*b*_ = 20 *k*_*B*_*T*. On the one hand, given that the Piezo dome’s area is ∼75% lipid membrane, this might not seem surprising. On the other hand, we wonder how the Piezo protein avoids adding stiffness to the Piezo dome—after all, the Piezo protein is an integral part of the dome’s structure. How Piezo creates the membrane dome by introducing intrinsic curvature without increasing stiffness is unknown. But the functional advantage afforded by these properties—intrinsic curvature without increased stiffness—can be understood when we consider their implications for sensing mechanical forces.

### Relating mechanics and structure to Piezo gating

Now we consider the implications of *R*_0_ ≈ 42 nm and *K*_*P*_ ≈ 18 *k*_*B*_*T* for the membrane dome model of mechanosensitive gating, using a thought experiment. To start, we embed a Piezo dome with energy 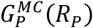 in Eq. 3 into an asymptotically planar free membrane that may be at a finite lateral tension *γ*. We refer to the Piezo dome plus its surrounding membrane as the ‘Piezo-membrane system.’ The shape energy of the Piezo-membrane system is given by

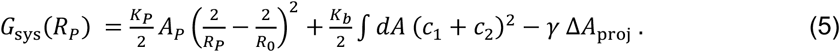

The first term in Eq. 5 is the Piezo dome mean curvature energy, Eq. 3. The second term is Eq. 1 for the free lipid bilayer membrane surrounding the Piezo dome. It captures, in analogy to the continuum elasticity theory describing the free membrane shape in Piezo vesicles, the membrane bending energy of Piezo’s membrane footprint (<u>accompanying paper</u>(8)). The third term is a work energy term that becomes important when a lateral tension is applied to the system(15). Δ*A*_proj_ < 0 thereby refers to the decrease in the projected, i.e., in-plane area of the Piezo-membrane system due to the curved shape of the Piezo dome and its membrane footprint, relative to the completely flat system configuration. For a given *R*_*P*_, we take Piezo’s membrane footprint to be in its minimum energy state obtained, similarly as for Piezo vesicles, by solving the corresponding Hamilton equations in an asymptotically flat, homogeneous bilayer membrane (see *Supplementary Information* section S2). As written, *G*_sys_(*R*_*P*_) is thus the shape energy of the Piezo-membrane system relative to the energy of a completely flat configuration of the system, i.e., with the Piezo dome and free membrane in a plane, without curvature. When *γ* = 0, the work energy term is equal to zero and so is the bending energy of Piezo’s membrane footprint, with the membrane footprint assuming a catenoidal shape in which *c*_1_ and *c*_2_ at every point are equal in magnitude and opposite in sign(8). In this case, Piezo’s membrane footprint exerts no force on the Piezo dome, and *G*_sys_(*R*_*P*_) attains its minimum at *R*_*P*_ = *R*_0_. Thus, when *γ* = 0, we have *G*_sys_ = 0. The left panel in Fig. 4*A* shows the shape of the Piezo-membrane system under nominal tension. The Piezo dome adopts its intrinsic radius of curvature, *R*_*P*_ ≈ 42 nm, and the surrounding free membrane forms a curved membrane footprint that smoothly meets the edge of the Piezo dome.

**Fig. 4:**
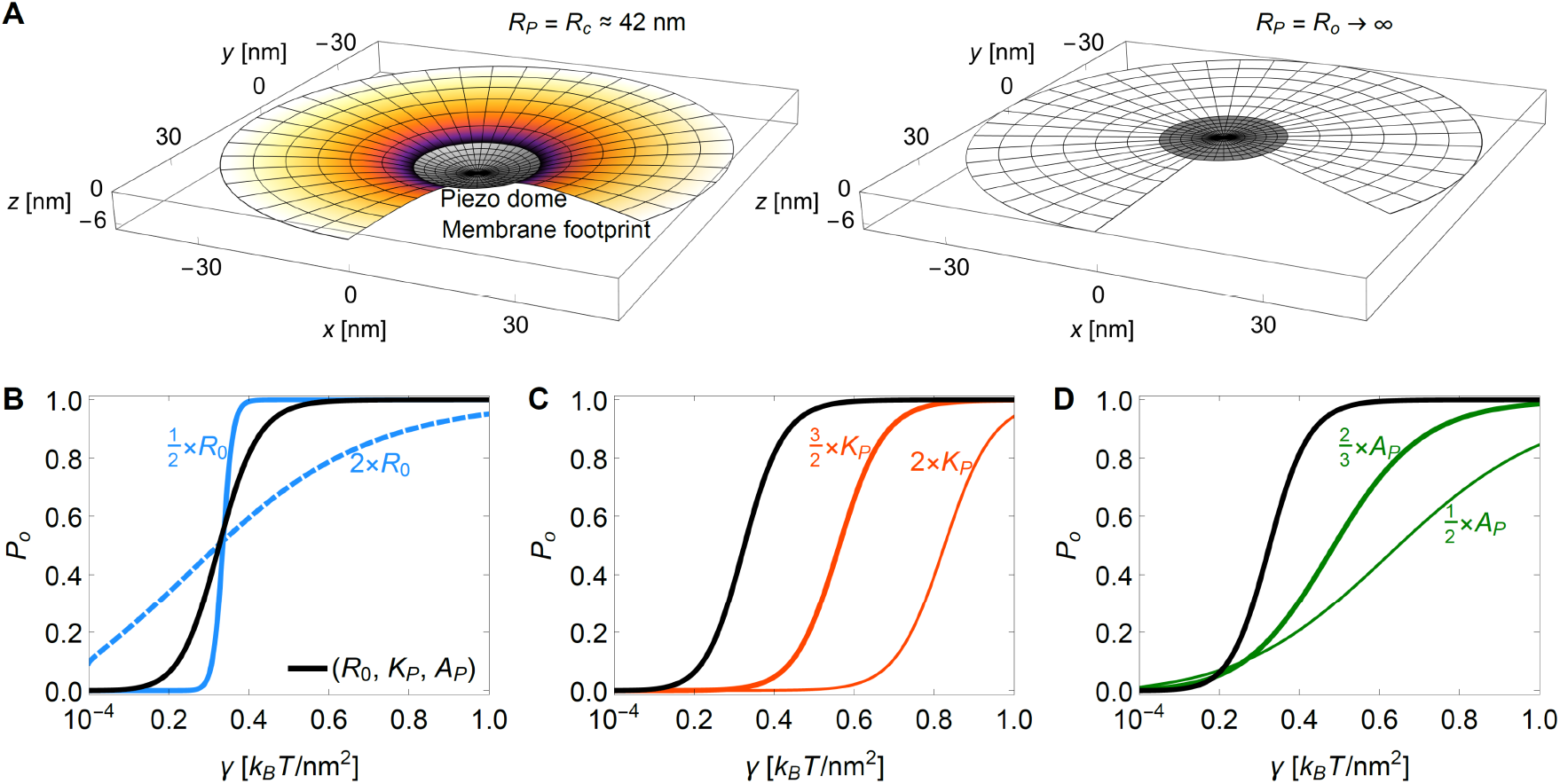
Mechanics of Piezo activation. (*A*) 3D plots of the Piezo dome and its (partial) membrane footprint in the limit of an infinitely large and asymptotically flat membrane. We assume here that the closed state of Piezo approximately corresponds to the intrinsic Piezo dome radius of curvature *R*_*P*_ = *R*_*c*_ ≈ 42 nm (left panel) and the open state to a flat Piezo dome shape with radius of curvature *R*_*P*_ = *R*_*o*_ → ∞ (right panel). We calculated the shape of the membrane footprint by minimizing the shape energy of the Piezo-membrane system, *G*_sys_(*R*_*P*_) in Eq. 5, at fixed *R*_*P*_. For the left panel we used a nominal lateral membrane tension 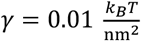. The membrane areas occupied by the Piezo dome, as well as by the membrane footprint, are identical in the left and right panels. (*B, C, D*) Piezo gating curves, *P*_*o*_(*γ*) in Eq. 6, for the elastic properties of the Piezo dome estimated in Fig. 3, *R*_0_ ≈ 42 nm and *K*_*P*_ ≈ 18 *k*_*B*_*T*, with the Piezo dome area *A*_*P*_ = 450 nm^2^ (black curves), and with (*B*) a modified intrinsic Piezo dome radius of curvature 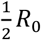 (blue solid curve) or 2*R*_0_ (blue dashed curve), (*C*) a modified Piezo dome bending rigidity 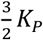 (thick red curve) or 2*K*_*P*_ (thin red curve), and (*D*) a modified Piezo dome area 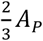 (thick green curve) or 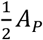 (thin green curve). For the predicted Piezo gating curve with *R*_0_ ≈ 42 nm, *K*_*P*_ ≈ 18 *k*_*B*_*T*, and *A*_*P*_ = 450 nm^2^ we have 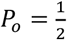 at the gating tension 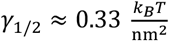 (black curves). In panel *B* we have 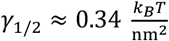 for 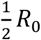 and 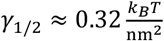 for 2*R*_0_, in panel *C* we have 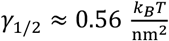 for 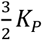 and 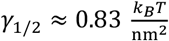 for 2*K*_*P*_, and in panel *D* we have 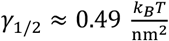 for 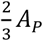 and 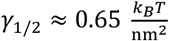 for 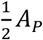.

When *γ* > 0, a flattening of the Piezo dome decreases the energetic cost of the work energy term in Eq. 5. At the same time, the energetic cost of curvature in the membrane footprint will also tend to flatten the Piezo dome. This is because, when *γ* > 0, the membrane footprint deviates from its catenoidal shape and the associated energy contribution must be positive, i.e., unfavorable, but less so if the membrane flattens out, reducing its curvature (8). Thus, when *γ* > 0, the second and third terms in Eq. 5 both favor a flatter, expanded configuration of the Piezo-membrane system, while the first term in Eq. 5 favors a shape of the Piezo dome with *R*_*P*_ = *R*_0_. As a result, the second and third terms in Eq. 5 yield a force on the Piezo dome to flatten it away from *R*_*P*_ = *R*_0_. This is how lateral membrane tension can alter the shape of the Piezo dome and its surrounding membrane (Fig. 4*A*, right panel).

As stated in the Introduction, we have proposed that the open conformation of Piezo is less curved. If we associate the closed Piezo channel with *R*_*P*_ = *R*_0_ (Fig. 4*A*, left panel) and the open channel with *R*_*P*_ → ∞ (Fig. 4*A*, right panel), i.e., a flat conformation, then applying the Boltzmann distribution equation to these two configurations of the Piezo-membrane system, we have

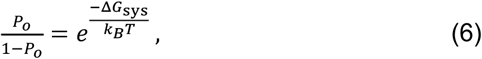

where *P*_*o*_ is the Piezo open probability and Δ*G*_sys_ is the difference in the shape energy of the Piezo-membrane system, Eq. 5, between the open and closed states of the Piezo channel. The black curve (see Figs. 4*B*-*D*) shows *P*_*o*_ as a function of *γ* according to Eqs. 5 and 6 for *A*_*P*_ = 450 nm^2^, *R*_0_ ≈ 42 nm, and *K*_*P*_ ≈ 18 *k*_*B*_*T*. We predict, with no free parameters, *P*_*o*_ = 0.5 at 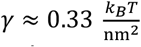, which is near published experimental values for the Piezo gating tension in cell membranes, with half activation around 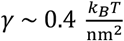 (16). The predicted and measured gating curves also show a comparable steepness.

Such a good correspondence between the predicted and measured tension-activation curves is surprising for several reasons. The assumption that the Piezo dome bending modulus, *K*_*P*_, for increasing the radius of curvature, i.e., flattening the dome, is the same as that for decreasing the radius of curvature, which we have measured, may be incorrect. Furthermore, Piezo is unlikely to be a 2-state channel and non-elastic (chemical) energy terms ought to contribute to the interaction between the Piezo protein and the lipid membrane, although it may be that these non-elastic energy terms do not change much between closed and opened conformations. With these assumptions understood, the elastic energy model of Piezo obtained here seems to approximate the functional behavior of the channel. And our motivation for this study is not to predict the activation curve, but to understand how the Piezo dome’s physical properties give rise to its mechanical force sensing ability. To this end, we systematically varied in our model Piezo’s intrinsic curvature, stiffness, and area to ask what effect these perturbations would have on its mechanosensing capability? The blue solid and dashed curves (Fig. 4*B*) show that Piezo’s steep, switch-like response to changes in membrane tension depends critically on the Piezo dome intrinsic radius of curvature, *R*_0_. Without intrinsic curvature, in its closed, resting state, Piezo would not be mechanosensitive. The red curves (Fig. 4*C*) show that Piezo’s ability to respond mechanically in the low-tension regime relies on the bending modulus of the Piezo dome being small. By small, we mean comparable to that of the pure lipid bilayer. If the value of *K*_*P*_ were doubled, for instance, then Piezo would not open until the membrane tension reached approximately one third of the lytic tension of a lipid bilayer, which is about 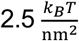 (12). Thus, intrinsic curvature of the Piezo dome with minimal stiffness appears to be an effective recipe for a switch-like conformational response in the low-tension regime.

The Piezo dome’s unusually large area is also important. The green curves (Fig. 4*D*) show that Piezo’s area influences both the steepness of its opening response and the tension range over which it opens. Thus, Piezo’s intrinsic curvature, membrane footprint, small bending modulus, and large area appear to be the key properties underlying Piezo’s ability to function as a highly responsive, tension-gated ion channel that operates in the low-tension regime.

## Discussion

Our analysis of Piezo vesicles is analogous to the problem of two connected springs, one whose force-displacement relationship is known and the other, unknown. By measuring the displacement of the dual spring system at mechanical equilibrium, because the forces between the two springs must be equal in magnitude and opposite in sign, the force-displacement relationship of the unknown spring can be deduced, without perturbing the system. In the present study, the vesicle free membrane represents one spring and the Piezo dome, the other. To designate the free membrane as known, we must know its force-displacement relationship. Thus, we examined in a companion paper whether the Helfrich energy equation can predict the shape of the free membrane bounded by the edge of the Piezo dome (<u>accompanying paper</u>). With no free parameters, we were able to accurately predict the shape of Piezo vesicles from the Helfrich energy equation. This justifies taking the Helfrich energy equation as a potential energy function for the free membrane, which means, through differentiation, we have the force. Thus, the free membrane serves as a known spring. Vesicles of different sizes permit the construction of a force-displacement relationship for the Piezo dome in unsupported, free standing lipid membranes.

To connect the force-displacement relationship of the Piezo dome to channel gating we model the Piezo dome as a spherical cap with intrinsic curvature *R*_0_, stiffness *K*_*P*_, and (known) area *A*_*P*_, whose deformation energy is governed by the dome’s mean curvature. Differentiation with respect to the Piezo dome radius of curvature, *R*_*P*_, yields a force function, Eq. 4, that conforms to the Piezo force curve data and gives estimates for *R*_0_ and *K*_*P*_. We conclude that Piezo in a planar, tensionless lipid bilayer membrane is curved, but less than in the detergent micelles used in cryo-EM studies. We also conclude that the Piezo dome exhibits low stiffness, comparable to that of the free lipid bilayer membrane.

The most interesting question in structural biology is not what does a molecule or collection of molecules look like, but why? Piezo is an extreme structural outlier amongst ion channels and membrane proteins in general. At its center, Piezo is a rather ordinary looking trimeric ion channel, but then each protomer extends radially a long, curved arm consisting of transmembrane helical units(6, 17, 18). Together, the three curved arms pucker the membrane to create a dome and surrounding membrane footprint that comprise a large area. The analysis we present argues that the structural features of large area and curvature are both important ingredients in the recipe for sensing lateral membrane tension. On top of these structural properties, the mechanical property of low stiffness is also important. To the question why regarding Piezo’s structure, we conclude that its large area, intrinsic curvature, and low stiffness are requirements for its ability to respond to membrane tension changes in the low-tension regime with high sensitivity. One could imagine in the evolution of its current form, that the Piezo channel began as an ‘ordinary’ ion channel, which became modified through natural selection to have extended arms to recruit a large dome area and membrane footprint, with a shape to produce intrinsic curvature, and with mechanical properties to ensure low bending stiffness.

Piezo’s solution to sensing membrane tension by an ion channel is not the only one that emerged in life. The mechanosensitive K^+^ channels TRAAK and TREK are small, wedge-shaped ion channels(19). Their shape probably produces a small membrane footprint, which would afford a *γ* Δ*A*_proj_ work energy term in the gating transition, but these channels’ small area is suboptimal. TRAAK and TREK open with a very weak dependence on membrane tension compared to Piezo (20). In another example, the large conductance bacterial mechanosensitive channel, MscL, opens with a strong dependence on membrane tension, like Piezo(21). MscL mainly produces its mechanical work term, *γ* Δ*A*_proj_, by direct, in-plane expansion of a disk-like arrangement of transmembrane helices that surround the pore(22). But there are two caveats. First, MscL functions as a pressure release valve in bacteria, opening a very wide pore in the face of osmotic shock. The large pore opening in MscL fulfills its role to release cytoplasmic content and to provide a strong tension dependence through a large Δ*A*_proj_, but such a large pore opening would be lethal to a eukaryotic cell. A eukaryotic cell must only open a narrow pore to mediate ion conduction, and thus pore opening alone will not produce a large Δ*A*_proj_. Second, MscL does not open until the membrane tension approaches lytic values(21). In other words, MscL does not open in the low-tension regime like Piezo. We would argue that the unique structure and elastic properties of Piezo reflect evolutionary adaptations to achieve strong tension dependent gating in the low-tension regime for an ion channel that opens a narrow pore.

In a previous study, HS-AFM was used to analyze shape changes in Piezo as a function of force applied by the AFM tip in imaging mode(9). Contrary to the present approach, the Piezo dome was thus perturbed by an external force applied from outside the membrane. The forces in that study were similar in magnitude to the forces estimated here, with both systems yielding estimates for the Piezo gating tension comparable to experimental values. The Piezo dome flattened and recoiled reversibly under the AFM tip, however, several observations and conclusions, especially regarding the Piezo dome geometry and force-displacement relationship, were different between the two studies. In the HS-AFM study, the low force intrinsic radius of curvature of the Piezo dome was about 15 nm rather than 40 nm. This difference alone makes a direct comparison of the two force curves impossible. One reason for the difference, we suspect, is that the HS-AFM study was carried out in lipid bilayer membranes made from POPE and POPG. In contrast to the lipids used in the present study, POPE and POPG produce membranes with a property sometimes called ‘intrinsic curvature’. These lipids do not favor formation of planar membrane sheets or spherical vesicles, have a tendency towards spontaneous curvature, and likely can organize around the curved Piezo channel, permitting it to be more curved than in POPC:DOPS:Cholesterol membranes(23). Another difference is that the membrane in the HS-AFM study was supported on a mica surface, whereas in the present study, the membrane is unsupported, more like a cellular membrane, with Piezo’s response to forces depending on its membrane footprint. Perhaps a useful conclusion to be reached by comparing these studies is that different lipid compositions and membrane environments in cells have the potential to regulate Piezo’s structure and thus its function, by biasing the Piezo dome towards curved or flattened states.

A recent structural study on Piezo in lipid vesicles(10) requires comment here to avoid misinterpretation of our findings and to ensure that the principles of membrane elasticity theory are correctly conveyed. In(10) the authors misuse the physical concepts ‘force’ and, in particular, ‘membrane tension,’ leading to an incorrect description of Piezo gating. They also specify the closed state radius of curvature of Piezo as 10 nm, the radius of curvature they observe in a spherical 10 nm radius vesicle. They use this radius of curvature to compare the predictions of the membrane dome model(6) to Piezo gating in a patch of membrane(16), which is not a 10 nm vesicle. The authors of(10) might have noticed in Figure 1 of their paper that in larger vesicles, Piezo’s radius of curvature is greater than 10 nm, as was also shown previously(9), and they ought to have wondered what it would be in a planar membrane. As we have shown, it is closer to 40 nm. This is because Piezo’s shape and function are inextricably tied to the geometry of the membrane through bending elastic forces, and emerge from Piezo-membrane interactions. This example raises the often-asked question in membrane protein structural biology, to what extent does the membrane, for example compared to a detergent micelle, alter a membrane protein’s structure? For Piezo, the answer is, a great extent. The beauty here is, membrane elasticity theory permits us to understand why, and this dependence of protein structure on the membrane is deeply rooted in Piezo’s mechanism of sensing mechanical force.

In summary, Piezo in planar lipid bilayer membranes is less curved than in detergent micelles, with a radius of curvature of about 40 nm at zero tension. But low threshold, sensitive mechanical gating properties are maintained nevertheless, owing to the creation of a membrane footprint surrounding Piezo. Realizing the biologically relevant shape and elastic properties of Piezo, which exploit the bending elastic properties of lipid bilayer membranes, is key to understanding its mechanosensory properties. The general concept of a dome model that utilizes *γ* Δ*A*_proj_ for Piezo’s mechanosensitive gating was hypothesized based on a structure in detergent micelles(6), but a more accurate description required an understanding of Piezo’s interaction with lipid bilayer membranes(8). In this and a companion paper (<u>accompanying paper</u>), we have quantified the bending elastic properties of Piezo inside a free-standing lipid bilayer membrane. Collectively, these findings tie Piezo’s unusual form and mechanical properties to its mechanosensing ability.

## Materials and Methods

For Figs. 1-3 we calculated the shape and elastic energy of the free membrane in Piezo vesicles as described in our companion paper (<u>accompanying paper</u>). We calculated the shape and elastic energy of Piezo’s membrane footprint in asymptotically planar membranes in Fig. 4 based on the theory developed in (8), using *Mathematica(24)*. A summary of this theoretical approach can be found in the *Supplementary Information* section S2 appended to this article.

## Data availability

The traced Piezo vesicle profiles underlying Figs. 1-4 are provided in the file ‘Traced Piezo vesicle profiles.txt’ available online with our companion paper (<u>accompanying paper</u>). Figures S1 and S2 can be found in the *Supplementary Information* section S1 appended to this article. The tomograms of Piezo vesicles are deposited to EMDataBbank database with the accession codes D_1000265122, D_100025123, D_1000265124, D_1000265125 and D_1000265126.

## Acknowledgments

This work was supported at USC by NSF Grant No. DMR-2051681 and by NSF Grant No. DMR-1554716 (to CAH) and at Rockefeller University by NIH Grant GM43949 (to RM). R.M. is an investigator of the Howard Hughes Medical Institute.

## Footnotes

1 Note that *A*_cap_ is not necessarily equal to the area of the Piezo dome, *A*_*P*_ ≈ 450 nm^2^. In particular, for the Piezo vesicles measured in our companion paper, we have 414 nm^2^ ≲ *A*_cap_ ≲ 471 nm^2^, with *A*_cap_ ≈ 421 nm^2^ in Fig. 2*A*.

## Notes

### Competing Interest Statement

The authors have declared no competing interest.

